# The heme precursor 5-aminolevulinic acid triggers the shutdown of the HemKR signalling system in *Leptospira*

**DOI:** 10.1101/2023.10.19.563136

**Authors:** Juan Andrés Imelio, Felipe Trajtenberg, Sonia Mondino, Leticia Zarantonelli, Iakov Vitrenko, Laure Lemée, Thomas Cokelaer, Mathieu Picardeau, Alejandro Buschiazzo

## Abstract

Heme and iron metabolic pathways are highly intertwined, both compounds being essential for key biological processes, yet becoming toxic if overabundant. Their concentrations are exquisitely regulated, including via dedicated two-component systems (TCSs) that sense signals and regulate adaptive responses. HemKR is a TCS involved in the control of heme metabolism in *Leptospira* spirochetes. However, the signals and molecular means by which HemKR is switched on/off, are still unknown. Moreover, a comprehensive list of HemKR-regulated genes, potentially overlapped with iron-responsive targets, is also missing. Here we show that 5-aminolevulinic acid (ALA), a committed porphyrin biosynthesis precursor, triggers the shutdown of the HemKR pathway by stimulating the phosphatase activity of HemK towards phosphorylated HemR. HemR dephosphorylation leads to differential expression of multiple genes, including of heme metabolism and transport systems. Furthermore, HemR inactivation brings about an iron-deficit tolerant phenotype, synergistically with iron-responsive signalling systems. Such tolerance could be vital during infection in pathogenic *Leptospira* species, which comprise a conserved HemKR TCS. In sum, HemKR responds to abundance of porphyrin metabolites by shutting down and controlling heme homeostasis, while also contributing to integrate the regulation of heme and iron metabolism in the *L. biflexa* spirochete model.

## INTRODUCTION

Bacteria detect intracellular and environmental stimuli through an array of sensory proteins that transmit the information, triggering physiologic adaptive responses. Such sensory/regulatory proteins constitute networks of signal transduction systems regulating numerous biologic processes (Imelio et al., 2021). Two-component systems (TCSs) are signal transduction systems (Jacob-Dubuisson et al., 2018; Francis & Porter, 2019), most often composed of a transmembrane sensory histidine-kinase (HK) and a cytosolic response regulator (RR). The presence/absence of a specific stimulus regulates the activation of the HK through ATP-mediated histidine autophosphorylation (Buschiazzo & Trajtenberg, 2019). The phosphorylated HK transfers its phosphoryl group to a conserved aspartate residue on its cognate RR (Gao et al., 2019), which can thereafter bind to DNA promoter sites, regulating the transcription of one or more genes. Most HKs are also capable to act as RR-specific phospho-aspartate phosphatases, critically contributing to keeping the pathways off when appropriate (Huynh et al., 2010; Gao & Stock, 2017; Buschiazzo & Trajtenberg, 2019). For such down-control function, some TCSs may also rely on dedicated auxiliary phosphatases, distinct from the sensory HK (Bourret & Silversmith, 2010; Parashar et al., 2011).

The Spirochetes phylum includes unique Gram-negative bacteria with spiral-shaped cells and endo-flagella confined within the periplasm (Wolgemuth et al., 2006). *Borrelia burgdorferi* (the causative agent of Lyme disease) and *Treponema pallidum* (syphilis), lack the enzymatic machinery to synthesize heme (Panek & O’Brian, 2002), with *Borrelia* evolving to use manganese instead of iron altogether (Posey & Gherardini, 2000). On the contrary, *Leptospira* spp. (including the causative agents of leptospirosis) need iron to survive (Louvel et al., 2006; Lo et al., 2010). Able to import heme from the milieu, *Leptospira* also synthesizes it *de novo* via the C5 pathway (Figure S1) (Guegan et al., 2003; Louvel et al., 2006). The metabolism of iron and heme are highly connected: heme is a tetrapyrrole that binds iron tightly within its porphyrin ring – an introduction that requires catalysis by a ferrochelatase–, allowing for heme’s essential functions in electron transfer, diatomic gas sensing/transport, and catalysis of diverse biochemical reactions (Dailey et al., 2017). Heme oxygenases on the other hand, degrade heme in an oxygen-dependent manner, rendering biliverdin, carbon monoxide and free iron (Frankenberg-Dinkel, 2004). Heme synthesis and catabolism thus depend on –and modify– the concentrations of iron in the cell. Intracellular iron excess is highly toxic, unleashing the production of harmful oxygen radicals through Fenton reactions. Excess of heme has also been reported to be toxic (Anzaldi & Skaar, 2010), even though the mechanisms by which heme kills bacteria remain unclear.

**Supporting Figure SI.**
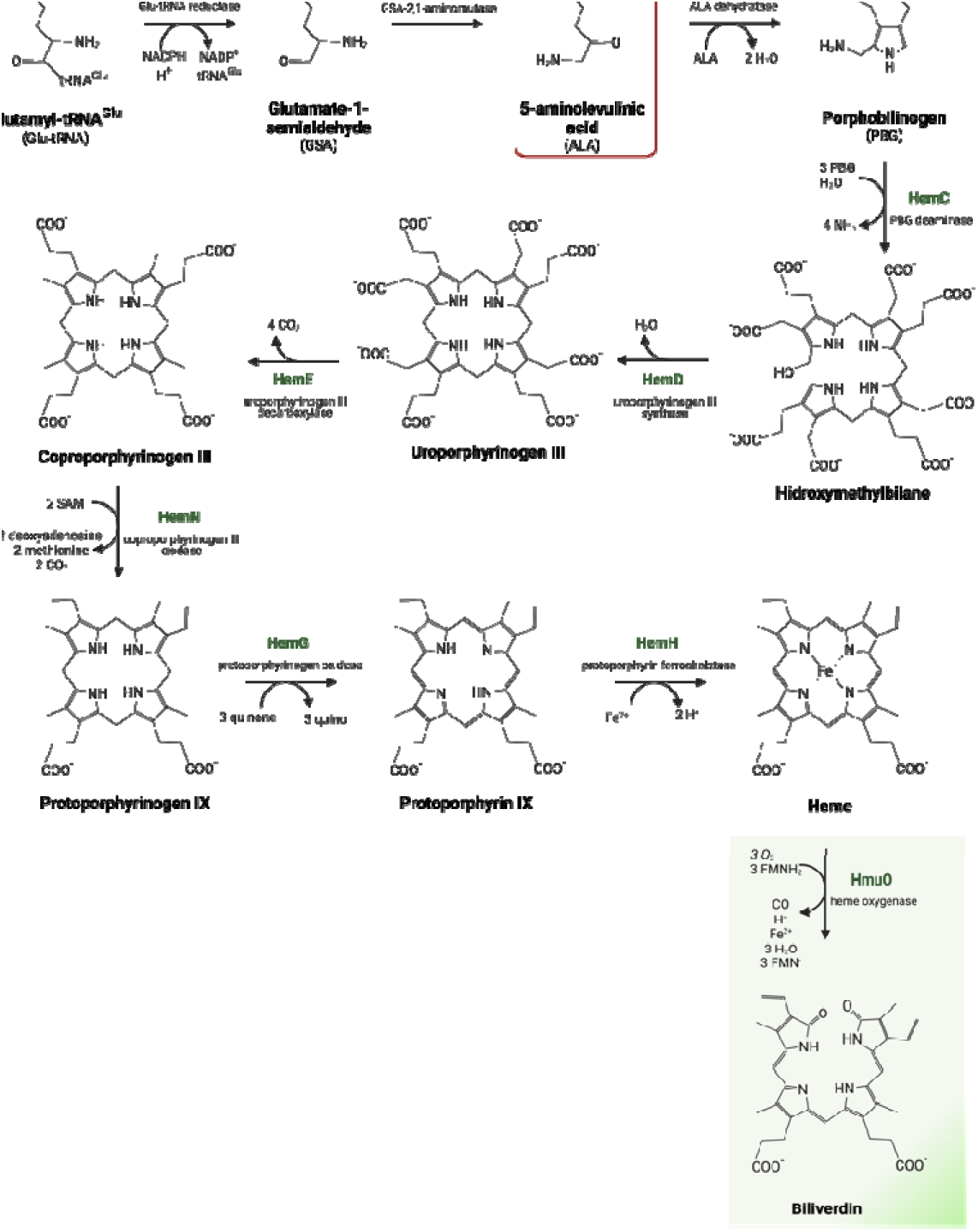
Heme biosynthesis In *Leptospira.* As In many other bacteria, *Leptospira* spp. possess all the genes coding for anabolic enzymes of the so called CS pathway for die *novo* heme biosynthesis. Starting from glutamyHRNA*^1^, as Initial precursor, the committed substrate 5-amlnolevullnlc add (ALA) Is generated (bowed In red). Four ALA moieties are needed to build the tetrapyrrole backbone of porphyrins Including heme. Note that HemC and HemD are found within a single blfunctional enzyme In *Leptospira* spp (Guegan et al„ 2003). The first step of heme degradation, catalysed by heme oxygenase, Is highlighted at the end of the pathway.

In bacteria, TCSs have been reported to sense iron (Steele et al., 2012). However, iron-signalling was shown to be mainly mediated via one-component systems, namely the ferric uptake regulator protein Fur and other iron-sensitive regulators such as DtxR/IdeR, RirA and Irr (Bradley et al., 2020). As for heme-responsive signalling, TCSs appear to play a dominant role in Gram positive bacteria (Keppel et al., 2020), whereas in Gram negatives, diverse examples point to extra-cytoplasmic function σ-factor signal transduction cascades, and direct/indirect sensing via iron-responsive Fur-like regulators (Kruger et al., 2022). In *Leptospira*, iron and heme are regulated in singular ways (Louvel et al., 2006), with no canonical Fur factors, and a distinct TCS, HemKR, controlling heme metabolism (Louvel et al., 2008). Using *Leptospira biflexa*, a saprophytic model that is similar to pathogenic species yet easier to manipulate (Picardeau, 2017), we have previously shown that HemR is phosphorylated by HemK, and that phospho-HemR (P∼HemR) binds to a *hem*-box (TGACA[N_6_]TGACA motif) present upstream of a few heme-metabolism genes (Morero et al., 2014). P∼HemR was proposed to be a transcriptional activator of genes encoding for the heme biosynthetic pathway, and a repressor of *hmuO*, which encodes the heme-degrading enzyme heme oxygenase (Morero et al., 2014). However, a more comprehensive examination of the HemR regulon was not available. Moreover, molecular signal(s) able to switch on/off the HemKR pathway, and the molecular mechanism by which such switching occurs in live cells, were still unknown.

In this work, we show that *L. biflexa* HemKR is shut down when the cells are exposed to extra-cellular 5-aminolevulinic acid (ALA), a committed porphyrin biosynthesis precursor. The signalling mechanism occurs by specific stimulation of HemK’s phosphatase activity over its cognate partner, P∼HemR. HemR dephosphorylation then leads to the differential expression of multiple genes, including of heme metabolism and transport systems. Lastly, we demonstrate that, synergistically with other iron-responsive systems, HemKR is involved in *L. biflexa* adaptation to iron starvation, ultimately conducing to bacterial iron-depletion tolerance.

## RESULTS

### The cytoplasmic catalytic region of HemK is trapped in the kinase-active state

A soluble construct of HemK (HemK_Δ80_) lacking the first 80 amino acids that span the periplasmic and trans-membrane regions of the native protein, possesses ATP-dependent autokinase activity (Figure 1a), confirming previous observations (Louvel et al., 2008). Further characterising the kinetics of the reaction, HemK_Δ80_ achieved 50% maximal phosphorylation in ∼60 min in the presence of excess ATP as phosphoryl donor (Figure 1a). P∼HemK_Δ80_ was also able to catalyse rapid phosphoryl-transfer to the phospho-receiver domain of HemR (HemR_REC_) (Figure 1b), with undetectable back-transfer (Figure 1c), similarly to what occurs in other HK families (Lima et al., 2023). Threonine 102 (T102) is predicted to play a key role in HemK’s phosphatase activity, as this is a conserved residue in this HK family (HisKA), engaged in the dephosphorylation of the RRs’ phospho-aspartate (Buschiazzo & Trajtenberg, 2019; Willett & Kirby, 2012). Neither the autophosphorylation nor the phosphoryl-transfer reaction kinetics were affected by substituting HemK’s T102 by an alanine (Figure 1a,b).

**Figure 1.**
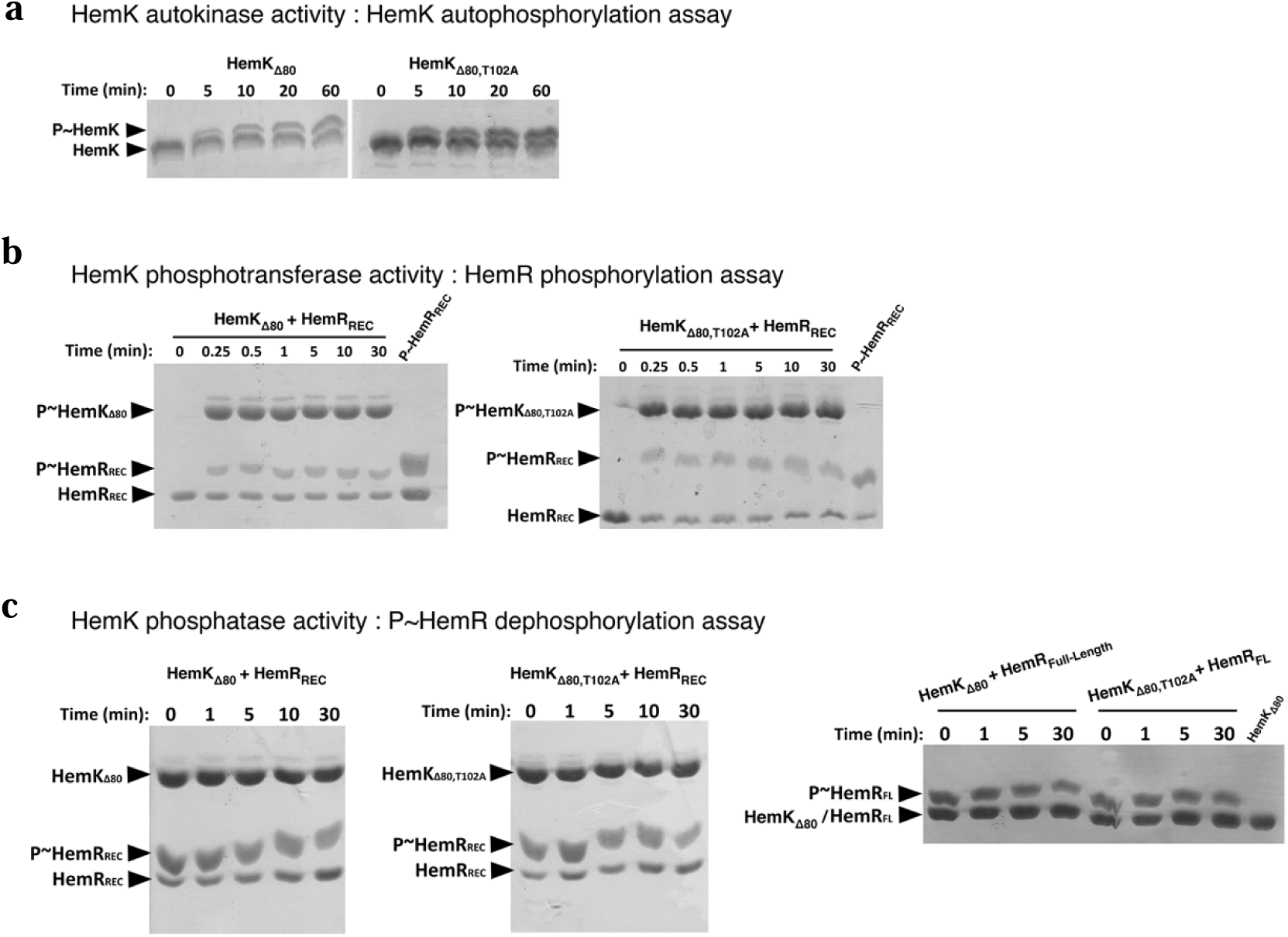
Autokinase, phosphotransferase and phosphatase activities catalysed by HemK. **(a)** Autophosphorylation of HemK_Δ80_ (left panel) and the phosphatase-null point-mutant HemK_Δ80,T102A_ (right), incubated for indicated times with 5mM ATP-Mg_2+_. **(b)** Phosphoryl-transfer kinetics from P∼HemK_Δ80_ (left) or P∼HemK_Δ80,T102A_ (right) to HemR_REC_, in the presence of 5mM ATP-Mg_2+_. **(c)** HemK-catalysed phosphatase activity on P∼HemR_REC_, using HemK_Δ80_ (left panel) or HemK_Δ80,T102A_ (centre). The dephosphorylation of the regulator was not detectable under any condition. 5mM ADP-Mg_2+_ was included in the reactions, as some HisKA His-kinases are known to require nucleotide to exert their phosphatase activity (indistinguishable results were obtained with no nucleotide added). The right-most panel reproduces the previous panels’ assays but using a full-length construct of HemR (HemR_FL_), ruling out the potential need of its DNA-binding domain to detect HemK phosphatase activity. The phosphorylated species of HemK and HemR are indicated (P∼HemK and P∼HemR, respectively) and suffer a mobility shift during electrophoresis due to the copolymerization of PhosTag into the polyacrylamide gels. Each experiment was performed in duplicate, representative Coomassie-stained gels are shown.

Even though most HisKA HKs exhibit specific phosphatase activity towards their cognate P∼RRs (Buschiazzo & Trajtenberg, 2019), HemK_Δ80_-dependent P∼HemR dephosphorylation was not detected *in vitro* (Figure 1c). Different conditions were assayed, especially considering known requirements of other HisKA kinases to fully catalyse phosphatase reactions. Among these, the inclusion of ADP (data not shown), and/or the use of full-length P∼HemR as substrate (Figure 1c), did not result in HemK_Δ80_ phosphatase enhancement.

Together with previous evidence exploring the effect of the key phosphorylatable histidine (H98) and aspartate (D53) residues on HemK and HemR, respectively (Louvel et al., 2008), we now show that HemK_Δ80_ specifically phosphorylates HemR in a unidirectional forward-transfer manner. Surprisingly, *in vitro* HemK_Δ80_ is not able to dephosphorylate P∼HemR, altogether suggesting that HemK_Δ80_ is intrinsically trapped in a kinase-active state, in need of its sensory/transmembrane portion to shift to a phosphatase-competent state under appropriate signalling conditions (as demonstrated below).

### The heme precursor 5-aminolevulinic acid stimulates HemK phosphatase activity towards P∼HemR

To discover signals sensed via the HemKR system, we asked whether the *L. biflexa* HemKR phosphorylation status was directly affected by exposition/deprivation of the cells to heme-synthesis building moieties (Figure 2). First attempts were done using heme itself. However, its toxicity at low micromolar concentrations, and the overwhelming global transcriptional effect we observed in *L. biflexa* cells exposed to <10µM hemin, hampered clearcut interpretations. Hence, a simpler scheme was followed using other heme-building blocks, such as 5-aminolevulinic acid (ALA) and iron. *L. biflexa* cells grown in EMJH medium until mid-exponential phase, were left untreated, or were supplemented with 300µM ALA, 35µM 2,2’-dipyridyl (DIP, a strong metal chelator) or 2mM FeSO_4_ (Figure 2). Bacterial whole protein extracts were then separated with PhosTag-SDS-PAGE, and HemR was revealed by Western blot using an anti-HemR polyclonal antibody (Figure 2a). We observed that in normal growth conditions (untreated) *L. biflexa* P∼HemR represents ∼12-25% of the total pool of HemR in the cells, and that this ratio did not change throughout the course of the experiment (Figure 2b). Excess or deficit of iron also did not produce detectable changes of HemR phosphorylation. However, exposure to ALA provoked a sustained depletion of P∼HemR, already observed at 1h of incubation (Figure 2a,b).

**Figure 2.**
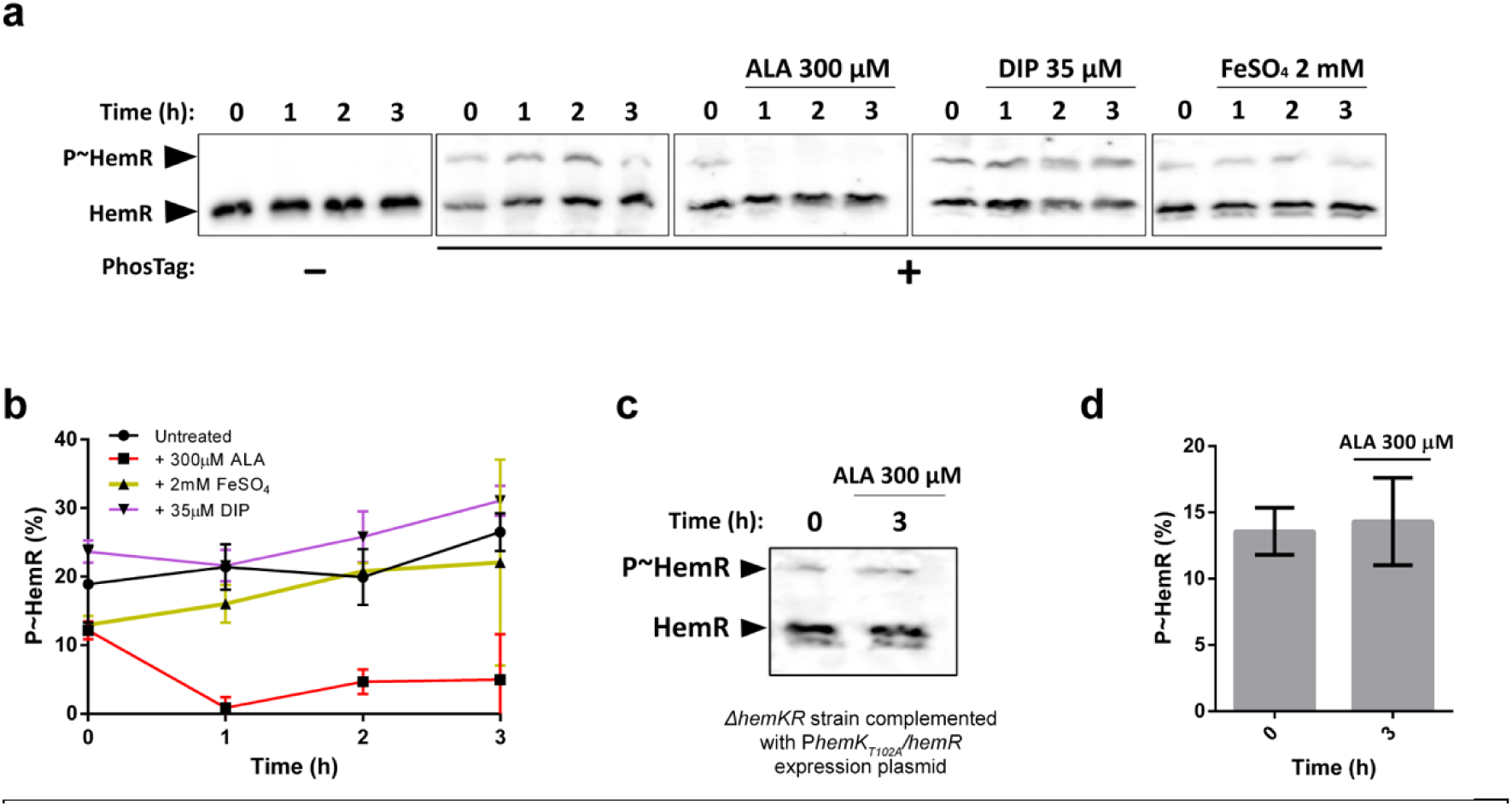
5-aminolevulinic acid (ALA), but not Fe^3+^, shuts down HemK-mediated phosphorylation of HemR *in vivo*. **(a)** PhosTag-SDS-PAGE of *L. biflexa wt* whole protein extracts, followed by Western blot using an anti-HemR antibody. Bacteria were left untreated or treated with 5-aminolevulinic acid (ALA), 2,2-dipyridyl (DIP), or an excess of iron (FeSO_4_), as indicated. Black arrows indicate the electrophoretic mobility position of phosphorylated and unphosphorylated HemR species. **(b)** Quantification of P∼HemR levels from panel (a) shown as percentage. Measurements were obtained from three biological replicates. **(c)** PhosTag-SDS-PAGE followed by Western blot as in panel (a). *L. biflexa* Δ*hemKR* mutant strain complemented with plasmid P*hemK_T102A_*/*hemR* was left untreated or treated with ALA for 3h. **(d)** Quantification of P∼HemR levels from panel (c), shown as percentage. Measurements were obtained from three biological replicates.

To uncover the molecular mechanism of this ALA-triggered effect, we hypothesized that HemK directly or indirectly senses ALA (or a downstream metabolite derived from ALA, including fully synthesized porphyrins such as heme), leading to the activation of its phosphatase activity towards P∼HemR. To test this hypothesis, we performed the same analysis using a *L. biflexa* Δ*hemKR* mutant strain (Louvel et al., 2008) complemented with plasmid P*hemK_T102A_*/*hemR*, encoding for HemK_T102A_ phosphatase-null mutant and *wt* HemR (Buschiazzo & Trajtenberg, 2019; Willett & Kirby, 2012) under the control of the operon’s native promoter. Notoriously, this strain exhibited comparable levels of HemR and P∼HemR as the *wt* strain, yet did not trigger P∼HemR dephosphorylation in response to ALA (Figure 2c,d).

Altogether, our results confirmed that HemK possesses phosphatase activity *in vivo*, leading to P∼HemR dephosphorylation in a signal-dependent manner. Indeed, ALA, but not iron, was shown to be a potential signal to shut down the HemKR pathway.

### HemKR-dependent gene regulation is consistent with a response effecting heme/iron homeostasis

Previous efforts to define the HemR regulon had pinpointed two targets, which code for enzymes involved in heme biosynthesis and degradation (Louvel et al., 2008; Morero et al., 2014). To further our understanding, we performed whole transcriptome analysis to identify HemKR-dependent differential gene expression (DGE) in *L. biflexa*, using ALA as an effective signal to shut down the pathway. We also used the wild-type strain (*wt*) and a double knockout with both *hemK* and *hemR* genes disrupted (Δ*hemKR*), to discriminate direct implications of HemKR in the ALA-triggered response (Table S1). *L. biflexa* single gene knockout strains (Δ*hemK* and Δ*hemR*) had been previously characterized as auxotrophic for heme, a deficiency that could also be rescued by adding ALA to the growth medium (Louvel et al., 2008). Intriguingly, *L. biflexa* Δ*hemKR* strain grew in the absence of ALA/heme, even though no evident differences could be identified among single and double knockout strains’ whole genome sequences (data not shown), other than the targeted gene disruption sites. The growth of the Δ*hemKR* strain was also not affected by washing thoroughly and passing the cells eight consecutive times in EMJH medium without ALA/heme supplementation, altogether confirming that this double knockout mutant is capable of *de novo* heme biosynthesis.

Transcriptomic analyses (Table S1) confirmed that transcription of *hemK* and *hemR* genes was indeed abolished in the mutant strain (see Experimental Procedures for further details). DGE analysis of ALA-treated *vs* untreated *wt Leptospira* cells, revealed a significant downregulation of 128 genes and upregulation of 51 (log_2_ fold change (FC) > |1|). This contrasted with the effect in the Δ*hemKR* mutant, which showed only 6 downregulated and 24 upregulated genes under the same experimental conditions (Figure 3a), consistent with HemKR being engaged in mediating the ALA-triggered response. Many ALA-responsive genes encode hypothetical proteins, but several key heme and iron metabolism-related genes were found to be differentially expressed in the *wt* strain (Figure 3a), with two representative targets (up- and down-regulated) further confirmed by qRT-PCR (Figure 3b, Table S2).

**Figure 3.**
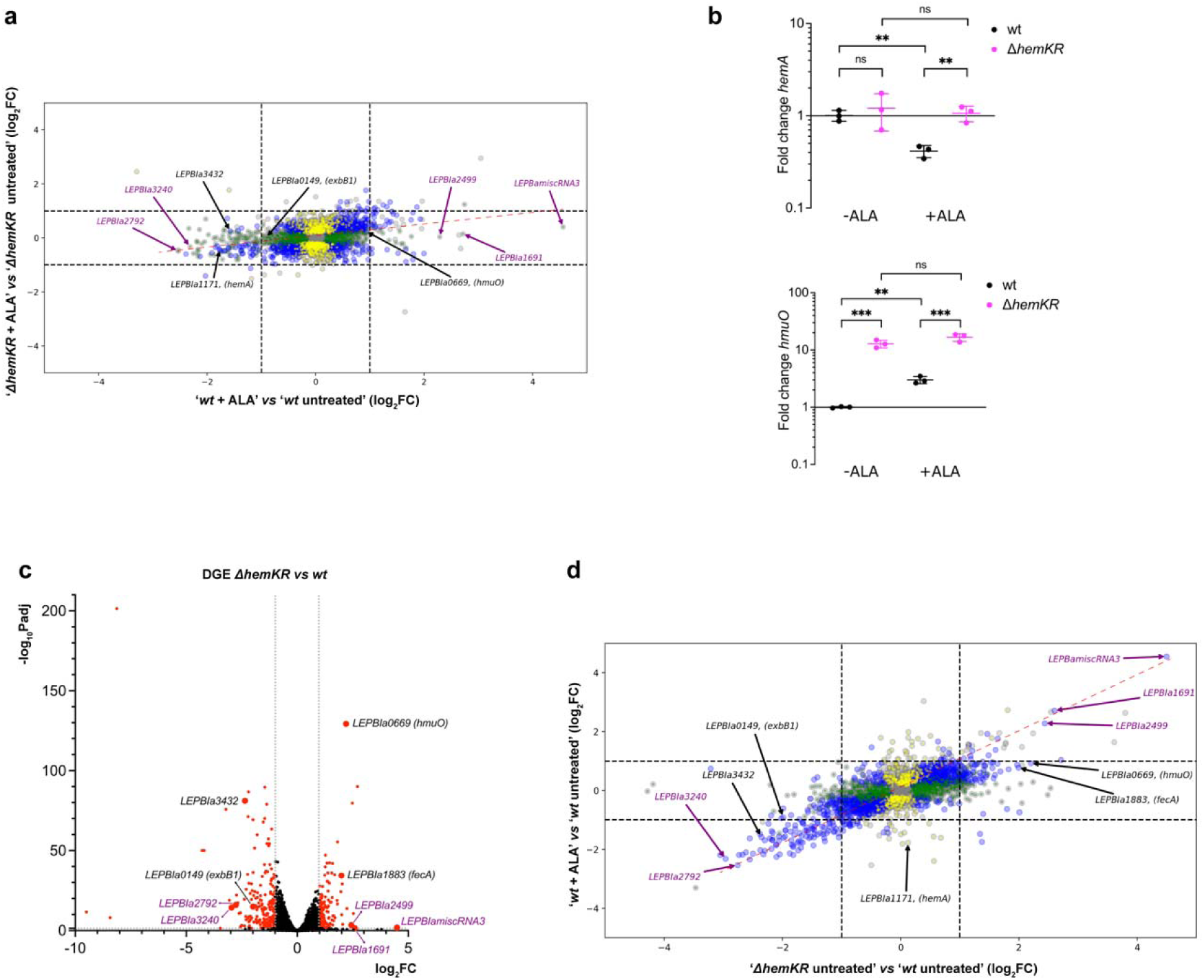
5-aminolevulinic acid (ALA) induces a largely HemKR-dependent transcriptional response. **(a)** Transcriptomic differential gene expression (DGE) analysis of Δ*hemKR* and *wt L. biflexa* strains, comparing cells that were treated with 300µM ALA vs untreated (see Table S1 for full data). The plot is a dual log_2_ fold change (log_2_FC), comparing the DGEs of Δ*hemKR vs wt* strains. Note the stronger regulation in the *wt* strain compared to the Δ*hemKR* mutant (flat pattern of log_2_FCs distribution; the red dashed line is the linear least squares regression curve with slope 0.1954 ± 0.011, and coefficient of determination R^2^=0.084). Blue dots represent genes with P-values (Padj) ≤ 0.05 in both strains; green dots, genes with Padj ≤ 0.05 only in *L. biflexa wt*; yellow dots, genes with Padj ≤ 0.05 only in *L. biflexa* Δ*hemKR*; grey dots, genes with Padj > 0.05. Black arrows label selected genes with functional annotation or clearcut homology to functionally characterized orthologs involved in heme/iron metabolism; purple arrows, correspond to hypothetical genes or non-protein-coding RNAs. **(b)** Confirmation of DGE effects triggered by ALA, performing qRT-PCR of two target genes, one repressed by ALA (*hemA*), and one induced (*hmuO*). Statistical significance of differences were calculated by Student’s *t* test using three replicates (ns=P>0.05; *=P≤0.05; **=P≤0.01;***=P≤0.001; ****=P≤0.0001). **(c)** Volcano representation of significance *vs* fold change analysing the transcriptomic DGE between *L. biflexa* Δ*hemKR vs wt* strains grown under standard EMJH culture conditions. Labelled genes are the same as in panel (a). **(d)** Dual log_2_FC plot comparing ‘*wt* ± ALA treatment’ vs ‘Δ*hemKR vs wt*’. Note the diagonal pattern of distribution (least squares regression in red dashed line, slope 0.4732 ± 0.009, and coefficient of determination R^2^=0.301), revealing a significant proportion of coincident differentially regulated genes in *wt*+ALA compared to Δ*hemKR*.

That the the dual log_2_FC plot of differential expression results in a horizontally flat distribution (Figure 3a), uncovers the stronger transcriptional response of *L. biflexa wt* to ALA treatment in comparison to the Δ*hemKR* mutant strain. Even though top-ranking differentially expressed genes included various targets related to heme metabolism –consistent with ALA triggering HemKR shutdown–, FC figures were modest. To obtain further confirmatory evidence, we then asked whether the removal of HemR, in the Δ*hemKR* strain, could be a relevant proxy of HemR dephosphorylation. Transcriptomic DGE was performed, comparing *L. biflexa* Δ*hemKR vs wt* strains grown in standard EMJH conditions (Figure 3c and Table S1). In general, abolishing HemKR resulted in similar effects to those visualized with the *wt* strain under ALA exposure, with the Δ*hemKR* knockout exhibiting even higher fold-change figures (Figures 3a and 3c). The dual log2FC plot in Fig 3d shows an overall linear relationship in the direction of differential expression suggesting a similar transcriptomic effect of the exposure to ALA and *hemKR* deletion, both leading to HemKR shutdown. The *hemA* operon was a notable exception, repressed in the *wt* after exposure to ALA while not significantly affected in the Δ*hemKR* mutant, it strongly suggests a composite control by different regulatory elements, which has been reported for this gene cluster in other bacteria (Aftab & Donegan, 2024).

Several of the differentially regulated genes, both in *wt*+ALA and in the Δ*hemKR* mutant, comprise *hem*-box motifs within their promoter regions, such as *hmuO* and *hemA,* the expression of which is indeed known to respond to HemR control (Morero et al., 2014). We now demonstrate that *exbB1/exbD1* and *LEPBIa3432* are also part of the HemR regulon. While the former had been previously anticipated (Morero et al., 2014), we have identified the consensus box **TGACA**GTACTG**TGACA** on the minus strand genomic coordinates 3550754-3550769, just upstream of *LEPBIa3430*/*LEPBIa3431*. We observed that these genes are indeed similarly downregulated as *LEPBIa3432*, altogether suggesting they form an operon. Several other ALA-responsive targets do not display easily discernible *hem*-box motifs, yet the majority correspond to genes encoding hypothetical proteins, justifying focused studies to unveil their regulatory mechanism and functional assignments.

In sum, ALA –and/or some derived metabolite along the porphyrin biosynthesis pathway– shuts down the HemKR system, anticipating heme synthesis inhibition (due to HemA decrease) and simultaneous stimulation of heme degradation (due to HmuO heme oxygenase’s surge).

### The phenotype of the ΔhemKR knockout strain revealed marked tolerance to iron deficit

Notoriously, *LEPBIa1883* lies among the most significantly upregulated genes in the *L. biflexa* Δ*hemKR* strain compared to *wt* (Figure 3c, 3d). This gene codes for the outer-membrane TonB-dependent ferric citrate receptor/transporter FecA. The expected effect would be for a larger pool of iron to become available in this mutant that lacks the HemKR system, convergent as well with the induced heme oxygenase that also liberates iron from heme. We thus conjectured that the Δ*hemKR* strain would be more tolerant to environmental iron deficit than the *wt*. To test this hypothesis, we assessed cell growth under increasing amounts of the iron-chelator, producing mild to severe iron deficiencies. While the *wt* strain displayed the expected sensitivity to iron deprivation, *L. biflexa* Δ*hemKR* was significantly more resistant (Figure 4).

**Figure 4.**
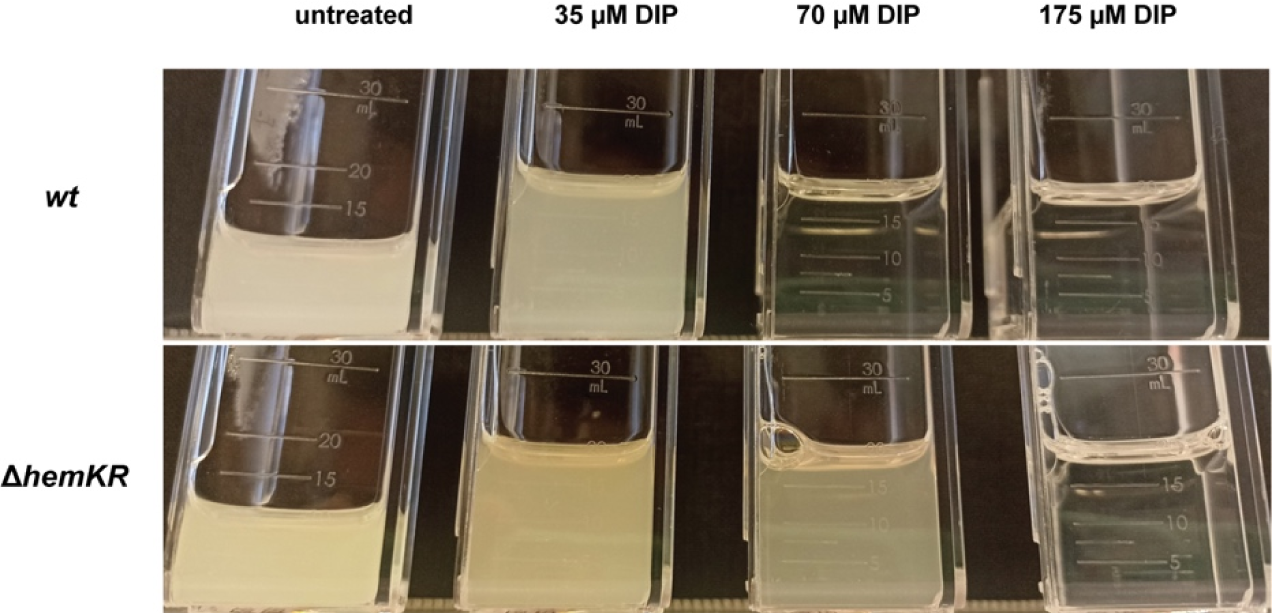
*L. biflexa* Δ*hemKR* strain is more tolerant to iron deficiency. *L. biflexa wt* and Δ*hemKR* were grown in EMJH medium with no supplemented iron. Cultures were left untreated or treated with increasing concentrations of the iron chelator 2,2’-dipyridyl (DIP), and incubated at 30°C with no agitation for 10 days. One representative experiment is shown from two biological

Complete depletion of iron can be achieved by further increasing the concentration of DIP (175 µM), at which point both *wt* and Δ*hemKR* cultures arrested growth, only to be reverted –in both strains–by adding the xenosiderophore deferoxamine (data not shown). All evidence considered, we confirmed that the *L. biflexa* Δ*hemKR* mutant, lacking the HemKR TCS, is indeed more tolerant to iron deficiency than the *wt* strain.

### Iron-deficiency triggers a transcriptional response that implicates a complex regulatory network beyond HemKR

The tolerance of *L. biflexa* Δ*hemKR* to iron deprivation suggested a crosstalk between heme- and iron-responsive regulatory pathways. We studied the systemic effect of iron on gene expression, in the *wt* and Δ*hemKR* strains. Iron-starving conditions were brought about by adding 35µM DIP to the culture, whereas iron excess was attempted by using 2mM FeSO_4_-supplemented medium (instead of 330 µM in standard EMJH). Unfortunately, iron overabundance hindered RNA-purification in the *wt* strain and could not be further analysed (Figure S2).

**Supporting Figure S2.**
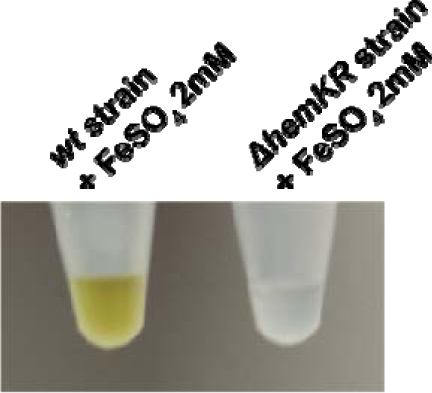
Samples, as labelled, were submitted to standard RNA extraction procedures in preparation for whole mRNA transcriptomic sequencing. For unknown reasons, *L. biflexa wt* strain –and not the Δ*hemKR* mutant– exhibited an abnormal behaviour if previously treated with excess iron: an insoluble yellow precipitate was formed during RNA extraction, which impaired RNA recovery prior to sequencing.

RNAseq analysis of *L. biflexa* under iron-restricted conditions revealed a strong regulatory response in both *wt* and Δ*hemKR* mutant (Figure 5a and Table S1). Largely consistent with reports studying *L. interrogans* (Lo et al., 2010) and other bacteria (Andrews et al., 2003) under similar stress, *L. biflexa wt* cells responded to Fe^3+^ deficit by downregulating 151 genes and upregulating 105 (considering DGEs of log_2_FC>|1|).

**Figure 5.**
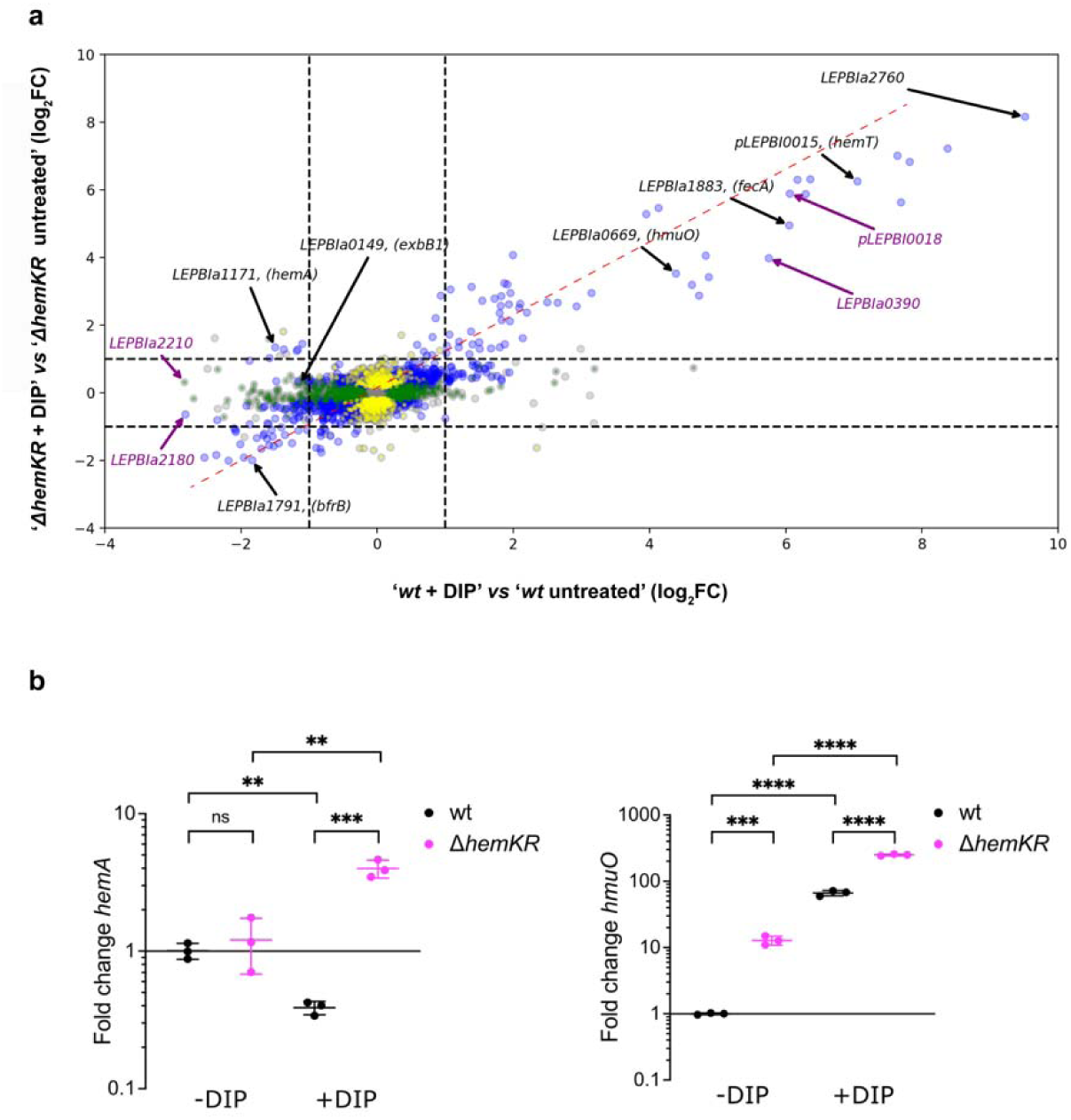
Iron deficit induces transcriptional regulation with mixed HemKR-dependent/independent responses. Differential gene expression analysis comparing *L. biflexa* strains treated *vs* untreated with 35 µM DIP for 3 h. **(a)** Dual log_2_ fold change (log_2_FC) plot, comparing differential gene expression of Δ*hemKR* and *wt* strains, treated with DIP *vs* untreated. Note the same global trend in differential expression between wt and mutant strains (diagonal pattern of log_2_FCs distribution; the red dashed line is the linear least squares regression curve, slope 0.5349 ± 0.009, coefficient of determination R^2^=0.456). Note however several off-diagonal genes suggesting synergistic effects of iron- and HemKR-dependent signalling. Colouring and labelling schemes as in Figure 3a. **(b)** Confirmation of DGE effects triggered by DIP, performing qRT-PCR of the same target genes as in Figure 3b (*hemA*, *hmuO*). Statistical significance of differences were calculated by Student’s t test using three replicates (ns=P>0.05; *=P≤0.05;

Although many of the iron-responsive genes code for hypothetical proteins, annotated genes linked to heme and/or iron metabolism were significantly affected, with representative targets further confirmed by qRT-PCR (Figure 5b, Table S2). Even though the pattern of the iron-deficit response was globally similar in *wt vs* Δ*hemKR* (upper right and lower left areas along the diagonal in Figure 5a), a significant number of genes displayed fold-change differences between both strains, strongly suggesting a synergistic overlap of HemKR and additional iron-responsive regulatory systems. For instance, a few genes encoding hypothetical proteins were over-expressed (*e.g. LEPBIa1691*, *LEPBIa0249*, among ∼25 others), whereas most were downshifted *(LEPBIa2210*, *LEPBIa2180*, among ∼100 others) only in the *wt* strain, and not in Δ*hemKR* (Figure 5a). Also of note, 17 tRNA-encoding genes (spanning tRNAs that bind to 13 different amino acids) were intriguingly overexpressed, only when the HemKR system is present. Importantly, the absence of the HemKR system, abolished the wild-type iron-sensitive behaviour of two operons that play key functions in heme homeostasis (*hemACBLENG* and *exbB1/exbD1*), which are repressed in the *wt* strain under iron-deficit (Figure 5a). Instead, in the Δ*hemKR* mutant *exbB1/exbD1* was no longer repressed and, more dramatically, the heme-biosynthesis operon *hemACBLENG* was induced ∼2.5-fold. This means that the Δ*hemKR* strain exhibited ∼8-fold increase of *hemACBLENG* expression compared to the *wt* under iron deprivation (Table S1), strikingly indicating a higher heme biosynthetic activity when the metal is restricted, that requires the HemKR system to be shut down.

All the evidence considered, a synergistic effect between HemKR and some other regulatory systems, facilitate an adaptive heme-homeostatic response when iron is scarce.

## Discussion

*Leptospira* cells require iron to live and are able to adapt in response to the metal’s availability (Lo et al., 2010; Louvel et al., 2006). *Leptospira* can also synthetise heme *de novo* (Guegan et al., 2003), a study that later led to the identification (Louvel et al., 2008) and characterization (Morero et al., 2014) of the two-component system HemKR as a regulator of heme metabolism. Two target genes, *hemA* and *hmuO*, were known to be respectively activated and repressed by P∼HemR (Morero et al., 2014), although more genes were suspected to be part of the regulon. Relevant questions remained concerning the signals that govern the phosphorylation status of the HemKR pathway in live cells, and the molecular mechanisms by which this TCS is controlled. Furthermore, whether HemKR could be implicated in coordinated responses linking heme and iron control, was not known.

We now report that 5-aminolevulinic acid (ALA) triggers the phosphatase activity of the sensory kinase HemK in live *L. biflexa* cells, promoting the dephosphorylation of its cognate response regulator HemR. The cytoplasmic, catalytically competent region of HemK, appears to be trapped in a constitutive kinase-on state, requiring the sensory/trans-membrane region of the protein to switch to a phosphatase configuration under the right signalling. This signal-driven dephosphorylation depends specifically on HemK’s phosphatase activity, eventually shutting down the HemKR pathway. Four residues away from the phosphorylatable histidine, HisKA family HKs (such as HemK) harbour an extremely conserved residue —a threonine or an asparagine—, critical to catalysing the hydrolytic dephosphorylation of their cognate P∼RR partner (Buschiazzo & Trajtenberg, 2019; Willett & Kirby, 2012). That position is occupied by threonine 102 in HemK, which after substitution by an alanine (T102A), proved that the phosphotransferase activity of HemK_T102A_ was not affected, yet it’s phosphatase activity was abolished, and with it, the capacity of HemK_T102A_ to sense and respond to ALA (Figure 2). Cues that turn off TCSs are well known, and play important biological roles such as in quorum sensing (Neiditch et al., 2006), chemotaxis (Parkinson et al., 2015) and virulence regulation (Lesne et al., 2018), among many others (Gao & Stock, 2017).

ALA is likely imported (Verkamp et al., 1993) in *L. biflexa* cells, and metabolized into heme and other porphyrins (Figure S1). Under the conditions of the experiments, we cannot distinguish whether ALA and/or any of the intermediary compounds along the porphyrin biosynthesis pathway, including heme itself, might be acting as specific signals for HemK. Heme’s toxicity on live *Leptospira* cells, and the extremely pleiotropic transcriptional response it triggers at sub-lethal concentrations, became challenging hurdles to address this question unequivocally. Nevertheless, ALA is a committed precursor of heme biosynthesis and thus well suited to act as a proxy of the cell’s heme-biosynthesis potential. Future biophysical/biochemical studies, using full-length trans-membrane HemK (*e.g.* with reconstituted proteoliposomes), are likely options to assess direct binding of candidate cues and subsequent kinase/phosphatase activity modulations.

Our transcriptomics approach to define a HemR regulon confirmed the two targets known to be regulated by phosphorylated HemR (P∼HemR), namely the *hemACBLENG* gene cluster and *hmuO* that encodes heme oxygenase. As expected from previous reports (Morero et al., 2014), *hemA* and *hmuO* were found to be down- and up-regulated respectively after exposure to ALA. In that study, Morero *et al*. had predicted that the transcription of the *exbB1/exbD1* gene cluster might also be controlled by P∼HemR, given the presence of a HemR-binding *hem*-box within its promoter region. Indeed, we now report that *exbB1/exbD1* is significantly repressed when the HemKR TCS is shut down by ALA (Figure 3a). ExbB1/ExbD1 is an inner-membrane subcomplex that acts in concert with TonB, to energize outer-membrane transporters engaged in iron/siderophore/heme import (Krewulak & Vogel, 2011). The shutdown of HemKR uncovered additional target genes of the regulon, such as *LEPBIa3432*, which has not been annotated, yet codes for a protein displaying clear homology to PhuR-like, TonB-dependent outer-membrane receptor/transporters (Richard et al., 2019). Together with ExbB1/ExbD1, a decrease of a putative heme receptor such as LEPBIa3432, concurrent with diminished HemA-dependent biosynthesis, anticipates a convergent down-control of heme concentrations. Added to heme oxygenase upregulation, when ALA is high (*i.e.* high heme-biosynthesis potential), cells maintain heme homeostasis which, if uncontrolled, would result in porphyrin overabundance.

Confirmation of the global expression effects of ALA-triggered HemKR shutdown, was obtained using the *L. biflexa* Δ*hemKR* mutant strain, which lacks HemR altogether. This mutant, a suitable proxy of HemR dephosphorylation, uncovered additional effects linking heme metabolism to iron availability. The overexpression of the iron transporter FecA, led to the hypothesis that the Δ*hemKR* mutant could be more tolerant to iron deficit. Indeed, a higher availability of intracellular iron due to increased transport (FecA) and augmented heme degradation (HmuO), could explain the greater tolerance of this mutant to iron deficit compared to *L. biflexa wt*. This scenario is also consistent with the strong overexpression of *LEPBIa1517*, which encodes a protein reliably predicted (Jumper & Hassabis, 2022) as a class 2 polyphosphate kinase PPK2 (Neville et al., 2022). PPK2s act as iron chelators, protecting bacterial cells from Fenton reaction toxicity (Beaufay et al., 2020) linked to higher iron concentrations.

Our analysis of the transcriptomic response to iron depletion was largely consistent with two previous studies on saprophytic and pathogenic *Leptospira* species (Louvel et al., 2005; Lo et al., 2010). Yet, we now uncovered a fairly large number of iron-responsive genes that were only affected in the *wt* strain and not in the Δ*hemKR* mutant (∼25 upregulated and >100 down-shifted), suggesting synergistic effects of HemKR and other signalling systems. Iron deprivation triggers a very strong regulatory response maximizing the entry of iron via heme and iron transporters, as well as mobilizing the intracellular iron pool by inducing heme demetallation – via heme oxygenase HmuO–, while repressing bacterioferritin-sequestered iron storage and heme biosynthesis (Figure 5). A direct role of the HemKR TCS is evidenced by the fact that the Δ*hemKR* mutant strain displays an altered expression of target genes, and more strikingly, inverts the effect on the heme-biosynthetic *hemA* operon. Additional transcription factors are yet to be identified to explain the overexpression of the *hemACBLENG* gene cluster in Δ*hemKR* when iron is lacking. The upregulation of both catabolic and biosynthetic heme enzymes in the DIP-treated Δ*hemKR* strain might be seen as an intriguing paradox. It is important to highlight that heme must be kept controlled when the heme-synthesis potential is high (such as under porphyrin precursors abundance). However, upon a deficit of environmental iron, cells must reallocate metabolic resources to release intracellular iron, while also securing enough heme to survive. It is tempting to speculate that this kind of reprogramming could account for the iron-deficit tolerance phenotype of *L. biflexa* Δ*hemKR* cells. Further investigations shall test this hypothesis, which predicts that HemKR’s ability to respond to ALA offers *Leptospira* cells a selective advantage under iron-limiting conditions, a typical scenario for pathogenic species during host infection (Ganz & Nemeth, 2015).

## EXPERIMENTAL PROCEDURES

### Bacterial strains and growth conditions

For targeted mutagenesis of the *hemK/hemR* operon in *Leptospira biflexa*, a suicide vector containing a kanamycin resistance cassette flanked by 600 bp of the 5’-end of *hemK* and 3’-end of *hemR* was introduced in *L. biflexa* serovar Patoc strain Patoc 1 by electroporation with a Biorad Gene Pulser Xcell, as previously described (Picardeau et al., 2001). Electroporated cells were plated on EMJH agar plates supplemented with 50 μg/ml kanamycin. Plates were incubated for 1 week at 30°C and kanamycin-resistant colonies were screened for double homologous recombination events by PCR. The *hemK*^-^/*hemR*^-^ double knockout mutant thus generated (named Δ*hemKR* in this study) was further confirmed by whole-genome sequencing. *Leptospira biflexa wt* and Δ*hemKR* strains were grown at 30°C in Ellinghausen-McCullough-Johnson-Harris (EMJH) liquid media (Goarant et al., 2020). When needed strains’ genotypes were confirmed by PCR with oligonucleotides HemK_Fw and HemR_Rev (Table S3). For growth measurements, *L. biflexa wt* and Δ*hemKR* were grown at 30°C in standard EMJH medium with no agitation. When cultures reached exponential phase (OD_420_ ∼0.25), a 1:100 inoculum of each strain was subcultured in EMJH medium without supplemented iron, in the presence of 0, 35, 70 or 175µM 2,2’-dipyridyl (DIP), at 30°C with no agitation until reaching stationary phase (10 days). Bacteria motility and viability were examined using a dark field microscope (AxioImager A2.LED, Zeiss).

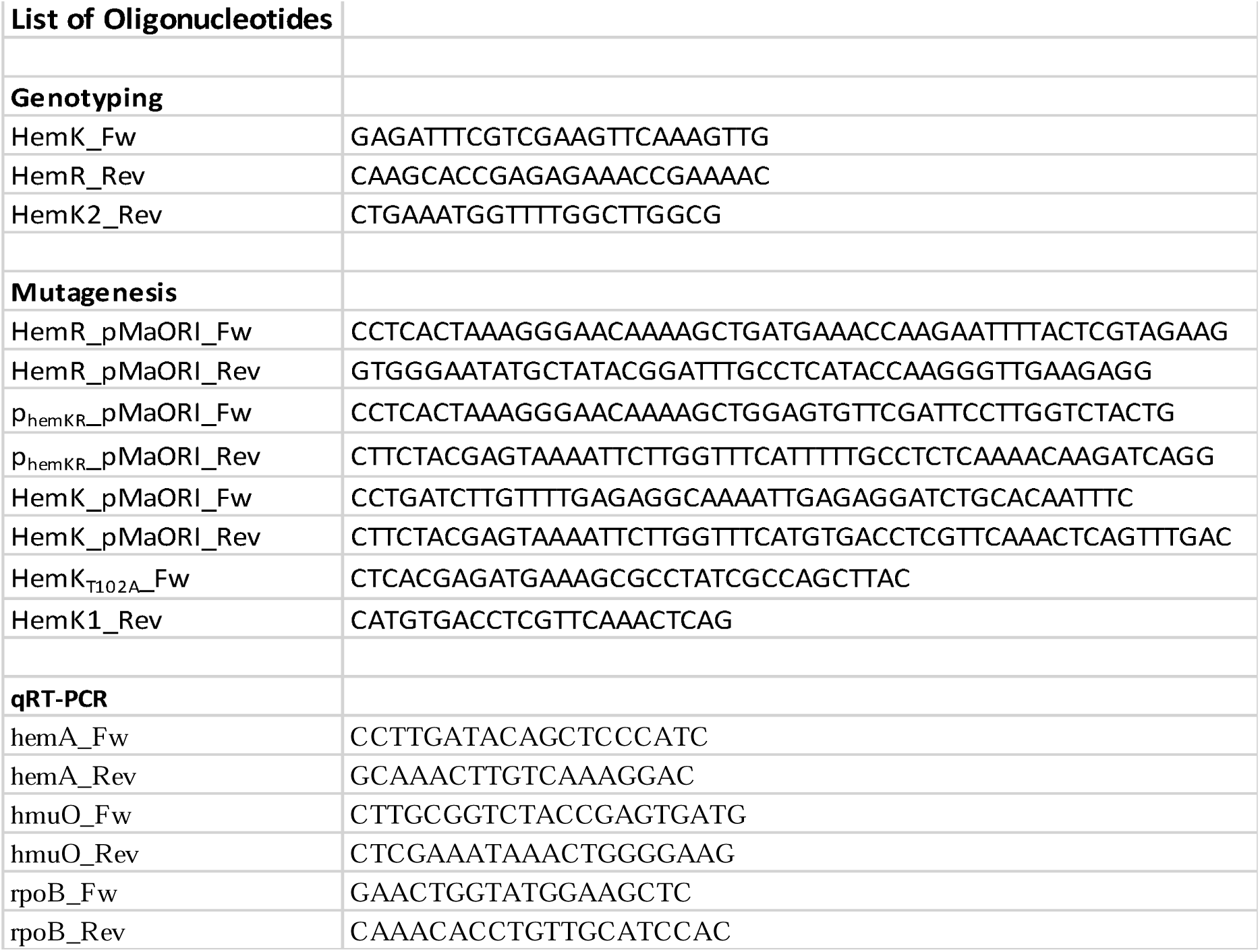
Supporting Table S3

The shuttle expression plasmid pMaORI (Spc^R^) (Pappas et al., 2015) was used to express mutated HemK-encoding gene constructs in *Leptospira*. Cloning and pMaORI manipulation/preparation was done in *E. coli* strain π1 (Δ*thyA*) cells (Demarre et al., 2005). pMaORI was transformed into *E. coli* strain β2163 (Δ*dapA*) (Demarre et al., 2005). *E. coli* strains π1 and β2163 were grown in LB containing 50 µg/mL spectinomycin, the former strain supplemented with 0.3 mM thymidine, whereas the latter with 0.3 mM diaminopimelic acid (DAP) (Demarre et al., 2005).

### Cloning of complementation plasmids

To complement *L. biflexa* Δ*hem*Κ*R* mutant, the wildtype *hemKR* allele harbouring its own promoter region (349 bp upstream *hemK*’s start codon), was amplified from *wt L. biflexa* using primers HemR_pMaORI_Fw, HemR_pMaORI_Rev, HemK_pMaORI_Fw, HemK_pMaORI_Rev, p_hemKR__pMaORI_Fw, and p_hemKR__pMaORI_Rev (Table S3). Site-directed mutagenesis was performed to generate a *hemK*_T102A_ mutant allele with primers HemK_T102A__Fw and HemK1_Rev (Table S3) and inserted into pMaORI (Pappas et al., 2015) by restriction-free cloning (van den Ent & Lowe, 2006). Correctly recombined pMaORI constructs were confirmed by BamHI/EcoRI digestion profile. Cloned pMaORI-*hemKR* and pMAORI-*hemK_T102A_R* plasmids were then transformed into *E. coli* strains π1 and β2163 for downstream conjugation experiments.

### Conjugation

*E. coli* strain β2163 cells harbouring the pMaORI-*hemKR* and pMaORI*-hemK_T102A_R* plasmids were co-incubated with *L. biflexa* cells in EMJH liquid media supplemented with 0.3 mM DAP (when required), until OD_420_ 0.3, according to standard procedures (Picardeau, 2008). Conjugants were incubated in solid EMJH agar plates supplemented with 50 µg/mL spectinomycin, at 30 °C for 7-10 days. Selected colonies were grown in liquid EMJH medium supplemented with 50 µg/mL spectinomycin, and the presence of the plasmid was confirmed by PCR with primers HemK_Fw and HemK2_Rev (Table S3).

### Expression and purification of recombinant proteins

pQE80L (Qiagen) plasmids harbouring HemK_Δ80_ (encoding a truncated version of the kinase, lacking the first 80 amino acids), HemK_Δ80,T102A_ (HemK_Δ80_ with threonine 102 substituted by alanine), HemR_REC_ (receiver domain only) and HemR_FL_ (full-length) ORFs, each with an encoded Tobacco Etch Virus (TEV) protease cleavage site for N-terminal His-tag removal, were transformed into *E. coli* Rosetta-gami(DE3) —for HemK— or TOP10F’ —for HemR— cells. Transformed Rosetta-gami(DE3) cells were grown with agitation (250 rpm) in 2xYT medium supplemented with 100 µg/mL ampicillin at 37°C until OD_600_ ∼ 1. Induction was carried with 1 mM IPTG at 30°C for 6 h. Transformed TOP10F’ cells were grown similarly, until OD_600_ ∼ 0.6. Induction was performed using 1 mM IPTG at 37°C for 3 h. Cells were harvested by centrifugation at 4000g, and pellets resuspended in 50 mM Tris pH 8, 500 mM NaCl, EDTA-free protease inhibitors (Roche) and 0.1 mg/mL DNase I (Sigma). Cells were disrupted with 1 mg/mL lysozyme followed by 1 cycle of freezing/thawing and high-pressure homogenisation (Emulsiflex-C5, Avestin).

Soluble fractions were obtained by centrifugation at 30000g for 30 min, and supernatants subjected to Ni^2+^ affinity chromatography (HisTrap, Cytiva) in buffer A (50 mM buffer Tris pH 8, 500 mM NaCl and 120 mM imidazole for recombinant HemK_Δ80_ and HemK_Δ80,T102A_ purification, or 20 mM imidazole for HemR_REC_ and HemR_FL_). Elution was performed with a gradient of buffer B (buffer A with added 750 mM imidazole for HemK_Δ80_ and HemK_Δ80,T102A_ purification, and 500 mM imidazole for HemR_REC_ and HemR_FL_). Eluted fractions were pooled and incubated overnight with TEV protease in dialysis buffer (50 mM Tris pH 8, 500 mM NaCl). Digested pools were subjected to a second Ni^2+^ column purification step to separate cleaved His-tag and TEV protease from the sample. All samples were finally subjected to size-exclusion chromatography (Superdex 16/60 75 prep column, Cytiva), previously equilibrated with 25 mM Tris pH 8 and 500 mM NaCl. The peak fractions were pooled, concentrated, and stored at -80°C until use.

### Autophosphorylation, phosphoryl-transfer and dephosphorylation catalysis assays

For autophosphorylation assays, 20 µM HemK_Δ80_ or HemK_Δ80,T102A_ were incubated with 5 mM ATP, 10 mM MgCl_2_. For phosphotransfer assays, 20 µM HemK_Δ80_ or HemK_Δ80,T102A_ previously incubated with 5 mM ATP, 10 mM MgCl_2_ for 60 min at room temperature (RT), were then incubated with equimolar concentrations of HemR_REC_ or HemR_FL_. For phosphatase assays, 600 µM HemR_REC_ or HemR_FL_ were first phosphorylated by incubating with 5 mM acetyl phosphate, 10 mM MgCl_2_, and then dimeric, phosphorylated HemR_REC_ or HemR_FL_, were purified from monomeric, non-phosphorylated forms, by size-exclusion chromatography (Superdex 10/300 75 prep, Cytiva). 20 µM of phosphorylated HemR_REC_ or HemR_FL_ were incubated with equimolar concentrations of HemK_Δ80_ or HemK_Δ80,T102A_. At defined time intervals, reactions were stopped with SDS buffer and 25 mM dithiothreitol (DTT) for 10 min at RT, followed by further incubation with 50 mM iodoacetamide for additional 10 min. All samples were then separated by PhosTag-SDS-PAGE, as described below. Gels were stained with Coomassie blue, scanned with a CanoScan Lide 110 (Canon) scanner, and bands quantified by densitometry using ImageJ (Schindelin et al., 2012).

### PhosTag-SDS-PAGE and Western blot of whole protein extracts

*L. biflexa* cultures were grown in 30 mL EMJH until OD_420_ ∼0.2. Prior to 5-aminolevulinic acid (ALA) or 2,2’-dipyridyl (DIP) incubation, an aliquot of 6.5 mL of untreated culture was harvested at 10000 g for 10 min at 4°C. When indicated, 300 µM ALA or 35 µM DIP was added to the cultures, and 6.5 ml aliquots were harvested at 1, 2 and 3 h post-incubation. Cell pellets were lysed resuspending with BugBuster and Lysonase according to the manufacturer’s protocol (Merck). Lysed cells were fractionated by centrifugation at 16000 g for 20 min at 4 °C, and an aliquot of the soluble fraction was resuspended in SDS buffer with 25 µM DTT and incubated 10 min at RT, followed by 50 µM iodoacetamide for extra 10 min at RT. Samples were run in SDS-PAGE 12% bis-acrylamide, co-polymerized with 100 µM PhosTag (Wako) and 200 µM ZnCl_2_. PhosTag in the presence of transition metals, shifts the electrophoretic mobility of phosphorylated proteins (Gao & Stock, 2018; Kinoshita et al., 2006), allowing to quantify P∼HemR/HemR ratios. Electrophoreses were run at <10 °C. Before Western blotting, PhosTag was eliminated by washing gels with SDS-PAGE running buffer and 2mM EDTA for 10 min, and then with no EDTA for an extra 10 min. Gels were blotted to nitrocellulose HyBond (Cytiva) membrane overnight with a TE 22 Mighty Small Transphor Tank Transfer Unit (Amersham) at 20 V.

A polyclonal monospecific anti-HemR antibody (αHemR) was produced in rabbits. Immunizations were done at Instituto Polo Tecnológico de Pando (Uruguay) respecting ethical guidelines for the use of animals in research. Briefly, 100 µg pure recombinant HemR (produced in *E. coli*, see (Morero et al., 2014) for construct and purification procedures) were injected subcutaneously with complete Freund’s adjuvant, followed by two boosters of 50 µg HemR in incomplete Freund’s at 14 and 28 days post-first immunization. Animals were bled for intermediate titration assays, and at ∼60 days post-immunization for sera separation, stored at - 20°C until use. Western blot membranes were pre-blocked with 3% bovine serum albumin in 0.1% PBS-Tween for 3.5 h, at RT and gentle agitation. αHemR was diluted 1/100 in blocking solution, and incubated with the membrane for 2 h at RT, with gentle agitation. Membranes were washed four times with 0.1% PBS-Tween 15 min/each, with strong agitation. Secondary anti-rabbit IgG (Sigma) antibody diluted 1/20000 in blocking solution was added to the membrane, with gentle agitation for 1 h RT. Four washing steps were repeated as before, and membranes were finally incubated with bioluminescent reagents (BM Chemiluminescence Western Blotting Kit, Roche) for 5 min. Protein bands were revealed and quantified using an ImageQuant^TM^ 800 (Cytiva) imaging system. Percentage of P∼HemR (%P∼HemR) was calculated by densitometry using ImageJ Fiji, subtracting background and subsequently quantifying total HemR signal (HemR_TOTAL_), which is the sum of the signal corresponding to P∼HemR plus the signal corresponding to non-phosphorylated HemR (HemR_NONP_) (HemR_TOTAL_ = P∼HemR + HemR_NONP_). Then, %P∼HemR = P∼HemR*100/HemR_TOTAL_.

### RNA extraction

*L. biflexa wt* and Δ*hemKR* mutant strains were grown in liquid EMJH media with no shaking at 30°C until mid-exponential phase (OD_420_ ∼0.2). Untreated cultures and cultures treated with FeSO_4_ 2 mM, δ-aminolevulinic acid (ALA) 300 µM, and 2,2’-dipyridyl (DIP) 35 µM were further incubated 3 h at 30°C with no shaking. RNA extraction was then performed as previously described (Zavala-Alvarado & Benaroudj, 2020).

### RNAseq

RNAseq libraries were prepared with 3 replicates for each condition, using a TruSeq Stranded mRNA library Preparation Kit (Illumina, USA) following the manufacturer’s protocol. The libraries were sequenced on an Illumina NextSeq2000 platform, generating single-end reads with an average read length of 107 bp. Quality control and data pre-processing were conducted to ensure robust downstream analyses. The number of reads was between 12-20M reads per sample, with an average phred score of 33. Transcription of the *hemK* and *hemR* genes were analysed in detail in the Δ*hemKR* mutant strain as a direct control of the knockout. *hemR* (*LEPBIa1422*) transcription was indeed abolished in all conditions. As for *hemK* (*LEPBIa1423*), standard differential gene expression (DGE) analyses did not detect a significant shift with respect to the *wt*; yet, a more detailed mapping on the genomic loci revealed that the reads corresponding to *hemK*, comprise only the initial 5’ fragment of the gene, upstream of the kanamycin-resistance cassette. Fold-change of individual mRNA species was an obvious figure to analyse DGE. The bioinformatic analyses of whole transcriptomes and differential gene expression were performed with Sequana (Cokelaer et al., 2017). Specifically, the RNAseq pipeline (v0.15.1) available at https://github.com/sequana/sequana_rnaseq was employed, built upon the Snakemake framework (Koster & Rahmann, 2012). To prepare the data, reads were trimmed for adapter sequences and low-quality bases using fastp software v0.20.1 (Chen et al., 2018) and were subsequently mapped to the *Leptospira* genome using bowtie2 (Langmead & Salzberg, 2012). The nucleotide sequence and annotation were downloaded from the MicroScope platform (Vallenet et al., 2020) using *L. biflexa* serovar Patoc Patoc 1 entry (corresponds to NCBI genome RefSeq assembly GCF_000017685.1). The count matrix was generated using FeatureCounts 2.0.0 (Liao et al., 2014), which assigned reads to relevant features based on the previously mentioned annotation. To identify differentially regulated genes, statistical analysis was performed with the DESeq2 library 1.30.0 (Love et al., 2014), and HTML reports were generated using the Sequana RNAseq pipeline. Key parameters for the statistical analysis encompassed significance, measured by Benjamini-Hochberg adjusted p-values with a false discovery rate (FDR) threshold of less than 0.05, as well as the effect size, quantified through fold-change calculations for each comparison.

### qRT-PCR

Transcription levels of target genes *hemA* (*LEPBIa1171*) and *hmuO* (*LEPBIa0669*) were evaluated in *L. biflexa wt* and Δ*hemKR* cells by quantitative real time PCR 9qRT-PCR). RNA extracted as described above, was used for cDNA synthesis using iScript cDNA Synthesis Kit (BioRad). PCR amplification was done on cDNAs using a QuantStudio 3 thermocycler (Applied Biosystems, Thermo Fisher Scientific), using the primers listed in Supporting Table S3, and followed in real time by using SsoFast EvaGreen Super-mix (BioRad), as follows: 15 min at 95°C; 40 x (0.25 min at 95°C, 1 min at 60°C); 0.25 min at 95°C; melting curves 1 min at 60°C; 0.25 min at 95°C with increment 0.1°C/s. The relative expression of target genes was calculated using the delta-delta Cq method and normalized against *rpoB* mRNA levels as described before (Morero et al., 2014).

## ACKNOWLEDGMENTS

JAI had PhD fellowships from ANII (POS_NAC_2016_1_ 129903) and CAP/Udelar. AB was supported by grants FCE_1_2017_1_136291 (ANII-Uruguay); and, together with MP, by the IMiZA Pasteur International Joint Research Unit (Inst Pasteur/Inst Pasteur Montevideo). Members of the Biomics Platform, C2RT, Institut Pasteur, Paris, France, were supported by France Génomique (ANR-10-INBS-09) and IBISA. We thank Horacio Botti for the lively discussions.

## REFERENCES

1. Aftab, H., and Donegan, R. K. (2024) Regulation of heme biosynthesis via the coproporphyrin dependent pathway in bacteria. Front Microbiol 15: 1345389.

2. Andrews, S. C., Robinson, A. K., and Rodriguez-Quinones, F. (2003) Bacterial iron homeostasis. FEMS Microbiol Rev 27: 215–237.

3. Anzaldi, L. L., and Skaar, E. P. (2010) Overcoming the heme paradox: heme toxicity and tolerance in bacterial pathogens. Infect Immun 78: 4977–4989.

4. Beaufay, F., Quarles, E., Franz, A., Katamanin, O., Wholey, W. Y., and Jakob, U. (2020) Polyphosphate Functions In Vivo as an Iron Chelator and Fenton Reaction Inhibitor. mBio 11.

5. Bourret, R. B., and Silversmith, R. E. (2010) Two-component signal transduction. Curr Opin Microbiol 13: 113–115.

6. Bradley, J. M., Svistunenko, D. A., Wilson, M. T., Hemmings, A. M., Moore, G. R., and Le Brun, N. E. (2020) Bacterial iron detoxification at the molecular level. J Biol Chem 295: 17602–17623.

7. Buschiazzo, A., and Trajtenberg, F. (2019) Two-Component Sensing and Regulation: How Do Histidine Kinases Talk with Response Regulators at the Molecular Level? Annual review of microbiology 73: 507–528.

8. Chen, S., Zhou, Y., Chen, Y., and Gu, J. (2018) fastp: an ultra-fast all-in-one FASTQ preprocessor. Bioinformatics 34: i884–i890.

9. Cokelaer, T., Desvillechabrol, D., Legendre, R., and Cardon, M. (2017) ‘Sequana’: a Set of Snakemake NGS pipeline. J Open Source Softw 2: 352.

10. Dailey, H. A., Dailey, T. A., Gerdes, S., Jahn, D., Jahn, M., O’Brian, M. R., and Warren, M. J. (2017) Prokaryotic Heme Biosynthesis: Multiple Pathways to a Common Essential Product. Microbiol Mol Biol Rev 81.

11. Demarre, G., Guerout, A. M., Matsumoto-Mashimo, C., Rowe-Magnus, D. A., Marliere, P., and Mazel, D. (2005) A new family of mobilizable suicide plasmids based on broad host range R388 plasmid (IncW) and RP4 plasmid (IncPalpha) conjugative machineries and their cognate Escherichia coli host strains. Res Microbiol 156: 245–255.

12. Francis, V. I., and Porter, S. L. (2019) Multikinase Networks: Two-Component Signaling Networks Integrating Multiple Stimuli. Annual review of microbiology 73: 199–223.

13. Frankenberg-Dinkel, N. (2004) Bacterial heme oxygenases. Antioxid Redox Signal 6: 825–834.

14. Ganz, T., and Nemeth, E. (2015) Iron homeostasis in host defence and inflammation. Nat Rev Immunol 15: 500–510.

15. Gao, R., Bouillet, S., and Stock, A. M. (2019) Structural Basis of Response Regulator Function. Annual review of microbiology 73: 175–197.

16. Gao, R., and Stock, A. M. (2017) Quantitative Kinetic Analyses of Shutting Off a Two-Component System. mBio 8: e00412–00417.

17. Gao, R., and Stock, A. M. (2018) Overcoming the Cost of Positive Autoregulation by Accelerating the Response with a Coupled Negative Feedback. Cell Rep 24: 3061–3071 e3066.

18. Goarant, C., Girault, D., Thibeaux, R., and Soupé-Gilbert, M.-E., (2020) Isolation and Culture of Leptospira from Clinical and Environmental Samples. In: Leptospira spp. Methods and Protocols N. Koizumi & M. Picardeau (eds). New York: Humana New York, pp. 1–9.

19. Guegan, R., Camadro, J. M., Saint Girons, I., and Picardeau, M. (2003) Leptospira spp. possess a complete haem biosynthetic pathway and are able to use exogenous haem sources. Mol Microbiol 49: 745–754.

20. Huynh, T. N., Noriega, C. E., and Stewart, V. (2010) Conserved mechanism for sensor phosphatase control of two-component signaling revealed in the nitrate sensor NarX. Proceedings of the National Academy of Sciences of the United States of America 107: 21140–21145.

21. Imelio, J. A., Trajtenberg, F., and Buschiazzo, A. (2021) Allostery and protein plasticity: the keystones for bacterial signaling and regulation. Biophys Rev 13: 943–953.

22. Jacob-Dubuisson, F., Mechaly, A., Betton, J. M., and Antoine, R. (2018) Structural insights into the signalling mechanisms of two-component systems. Nat Rev Microbiol 16: 585–593.

23. Jumper, J., and Hassabis, D. (2022) Protein structure predictions to atomic accuracy with AlphaFold. Nature methods 19: 11–12.

24. Keppel, M., Hunnefeld, M., Filipchyk, A., Viets, U., Davoudi, C. F., Kruger, A., Frunzke, J. (2020) HrrSA orchestrates a systemic response to heme and determines prioritization of terminal cytochrome oxidase expression. Nucleic Acids Res 48: 6547–6562.

25. Kinoshita, E., Kinoshita-Kikuta, E., Takiyama, K., and Koike, T. (2006) Phosphate-binding tag, a new tool to visualize phosphorylated proteins. Mol Cell Proteomics 5: 749–757.

26. Koster, J., and Rahmann, S. (2012) Snakemake--a scalable bioinformatics workflow engine. Bioinformatics 28: 2520–2522.

27. Krewulak, K. D., and Vogel, H. J. (2011) TonB or not TonB: is that the question? Biochem Cell Biol 89: 87–97.

28. Kruger, A., Keppel, M., Sharma, V., and Frunzke, J. (2022) The diversity of heme sensor systems heme-responsive transcriptional regulation mediated by transient heme protein interactions. FEMS Microbiol Rev 46.

29. Langmead, B., and Salzberg, S. L. (2012) Fast gapped-read alignment with Bowtie 2. Nature methods 9: 357–359.

30. Lesne, E., Dupre, E., Lensink, M. F., Locht, C., Antoine, R., and Jacob-Dubuisson, F. (2018) Coiled-Coil Antagonism Regulates Activity of Venus Flytrap-Domain-Containing Sensor Kinases of the BvgS Family. mBio 9: e02052–02017.

31. Liao, Y., Smyth, G. K., and Shi, W. (2014) featureCounts: an efficient general purpose program for assigning sequence reads to genomic features. Bioinformatics 30: 923–930.

32. Lima, S., Blanco, J., Olivieri, F., Imelio, J. A., Nieves, M., Carrion, F., Trajtenberg, F. (2023) An allosteric switch ensures efficient unidirectional information transmission by the histidine kinase DesK from Bacillus subtilis. Sci Signal 16: eabo7588.

33. Lo, M., Murray, G. L., Khoo, C. A., Haake, D. A., Zuerner, R. L., and Adler, B. (2010) Transcriptional response of Leptospira interrogans to iron limitation and characterization of a PerR homolog. Infect Immun 78: 4850–4859.

34. Louvel, H., Betton, J. M., and Picardeau, M. (2008) Heme rescues a two-component system Leptospira biflexa mutant. BMC Microbiol 8 25.

35. Louvel, H., Bommezzadri, S., Zidane, N., Boursaux-Eude, C., Creno, S., Magnier, A., Picardeau, M. (2006) Comparative and functional genomic analyses of iron transport and regulation in Leptospira spp. Journal of bacteriology 188: 7893–7904.

36. Louvel, H., Saint Girons, I., and Picardeau, M. (2005) Isolation and characterization of FecA- and FeoB-mediated iron acquisition systems of the spirochete Leptospira biflexa by random insertional mutagenesis. Journal of bacteriology 187: 3249–3254.

37. Love, M. I., Huber, W., and Anders, S. (2014) Moderated estimation of fold change and dispersion for RNA-seq data with DESeq2. Genome Biol 15 550.

38. Morero, N. R., Botti, H., Nitta, K. R., Carrion, F., Obal, G., Picardeau, M., and Buschiazzo, A. (2014) HemR is an OmpR/PhoB-like response regulator from Leptospira, which simultaneously effects transcriptional activation and repression of key haem metabolism genes. Mol Microbiol 94: 340–352.

39. Neiditch, M. B., Federle, M. J., Pompeani, A. J., Kelly, R. C., Swem, D. L., Jeffrey, P. D., Hughson, F. M. (2006) Ligand-induced asymmetry in histidine sensor kinase complex regulates quorum sensing. Cell 126: 1095–1108.

40. Neville, N., Roberge, N., and Jia, Z. (2022) Polyphosphate Kinase 2 (PPK2) Enzymes: Structure, Function, and Roles in Bacterial Physiology and Virulence. Int J Mol Sci 23.

41. Panek, H., and O’Brian, M. R. (2002) A whole genome view of prokaryotic haem biosynthesis. Microbiology (Reading) 148: 2273–2282.

42. Pappas, C. J., Benaroudj, N., and Picardeau, M. (2015) A replicative plasmid vector allows efficient complementation of pathogenic Leptospira strains. Appl Environ Microbiol 81: 3176–3181.

43. Parashar, V., Mirouze, N., Dubnau, D. A., and Neiditch, M. B. (2011) Structural basis of response regulator dephosphorylation by Rap phosphatases. PLoS biology 9: e1000589.

44. Parkinson, J. S., Hazelbauer, G. L., and Falke, J. J. (2015) Signaling and sensory adaptation in Escherichia coli chemoreceptors: 2015 update. Trends Microbiol 23: 257–266.

45. Picardeau, M. (2008) Conjugative transfer between Escherichia coli and Leptospira spp. as a new genetic tool. Appl Environ Microbiol 74: 319–322.

46. Picardeau, M. (2017) Virulence of the zoonotic agent of leptospirosis: still terra incognita? Nat Rev Microbiol 15: 297–307.

47. Picardeau, M., Brenot, A., and Saint Girons, I. (2001) First evidence for gene replacement in Leptospira spp. Inactivation of L. biflexa flaB results in non-motile mutants deficient in endoflagella. Mol Microbiol 40: 189–199.

48. Posey, J. E., and Gherardini, F. C. (2000) Lack of a role for iron in the Lyme disease pathogen. Science 288: 1651–1653.

49. Richard, K. L., Kelley, B. R., and Johnson, J. G. (2019) Heme Uptake and Utilization by Gram-Negative Bacterial Pathogens. Front Cell Infect Microbiol 9: 81.

50. Schindelin, J., Arganda-Carreras, I., Frise, E., Kaynig, V., Longair, M., Pietzsch, T., Cardona, A. (2012) Fiji: an open-source platform for biological-image analysis. Nature methods 9: 676–682.

51. Steele, K. H., O’Connor, L. H., Burpo, N., Kohler, K., and Johnston, J. W. (2012) Characterization of a ferrous iron-responsive two-component system in nontypeable Haemophilus influenzae. Journal of bacteriology 194: 6162–6173.

52. Vallenet, D., Calteau, A., Dubois, M., Amours, P., Bazin, A., Beuvin, M., Medigue, C. (2020) MicroScope: an integrated platform for the annotation and exploration of microbial gene functions through genomic, pangenomic and metabolic comparative analysis. Nucleic Acids Res 48: D579–D589.

53. van den Ent, F., and Lowe, J. (2006) RF cloning: a restriction-free method for inserting target genes into plasmids. J Biochem Biophys Methods 67: 67–74.

54. Verkamp, E., Backman, V. M., Bjornsson, J. M., Soll, D., and Eggertsson, G. (1993) The periplasmic dipeptide permease system transports 5-aminolevulinic acid in Escherichia coli. Journal of bacteriology 175: 1452–1456.

55. Willett, J. W., and Kirby, J. R. (2012) Genetic and biochemical dissection of a HisKA domain identifies residues required exclusively for kinase and phosphatase activities. PLoS Genet 8: e1003084.

56. Wolgemuth, C. W., Charon, N. W., Goldstein, S. F., and Goldstein, R. E. (2006) The flagellar cytoskeleton of the spirochetes. J Mol Microbiol Biotechnol 11: 221–227.

57. Zavala-Alvarado, C., and Benaroudj, N., (2020) The Single-Step Method of RNA Purification Applied to Leptospira. In: Leptospira spp. Methods and Protocols N. Koizumi & M. Picardeau (eds). New York: Humana New York, pp. 41–51.

