## Supplementary figures and images for "The heme precursor 5-aminolevulinic acid triggers the shutdown of the HemKR signalling system in *Leptospira*"

### Supplemental Figure S1

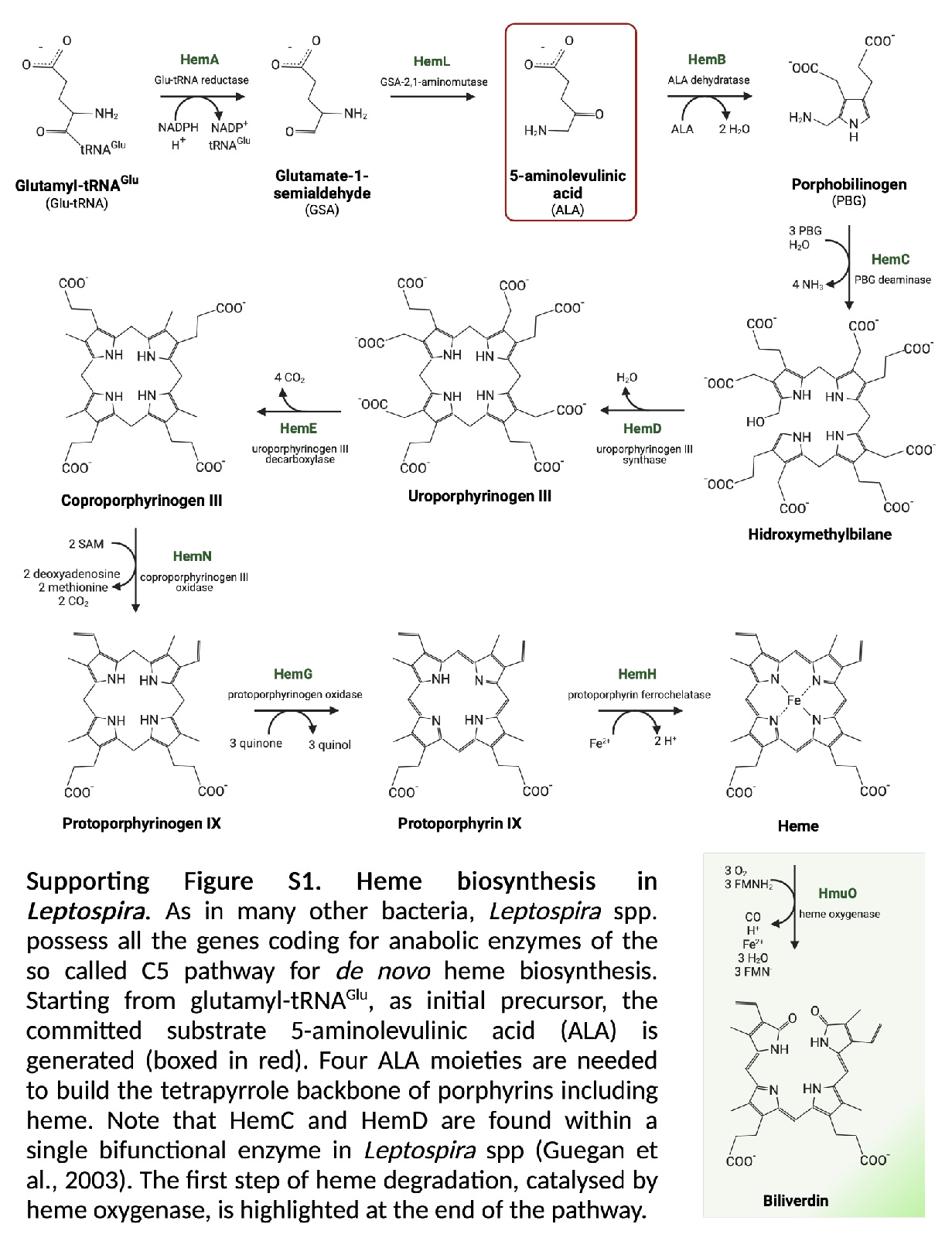

### Supplemental Figure S2

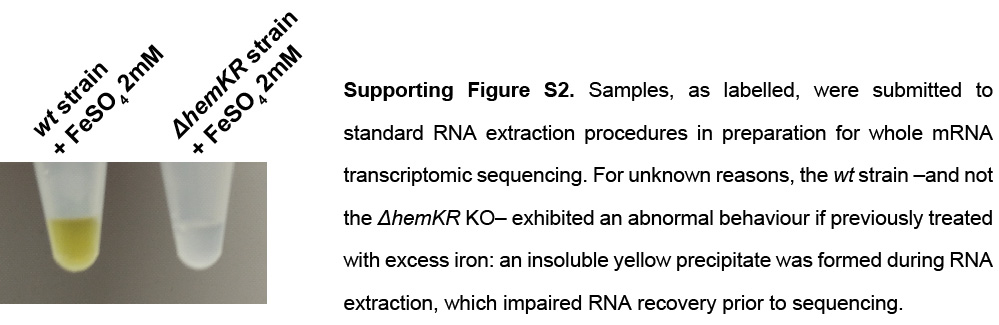
